# Mapping molecular HLA typing data to UNOS antigen equivalents for improved virtual crossmatch

**DOI:** 10.1101/328914

**Authors:** Navchetan Kaur, Evan P. Kransdorf, Marcelo J Pando, Martin Maiers, Bryan Ray, Jar-How Lee, Peter Lalli, Cathi L Murphey, Robert A Bray, Loren Gragert

**Author notes:** Corresponding author: Loren Gragert, 1430 Tulane Ave, P.O. Box 8679, New Orleans, LA 70112.

## Abstract

**Background:** Virtual crossmatch utilizes HLA typing and antibody screen assay data as a part of organ offers in deceased donor allocation systems. Histocompatibility labs must convert molecular HLA typings to antigen equivalencies for entry into the United Network for Organ Sharing (UNOS) UNet system. While an Organ Procurement and Transplantation Network (OPTN) policy document provides general guidelines for conversion, the process is complex because no antigen mapping table is available. We present a UNOS antigen equivalency table for all IMGT/HLA alleles at the A, B, C, DRB1, DRB3/4/5, DQA1, and DQB1 loci.

**Methods:** An automated script was developed to generate a UNOS antigen equivalency table. Data sources used in the conversion algorithm included the World Marrow Donor Association(WMDA) antigen table, the HLA Dictionary, and UNOS-provided tables. To validate antigen mappings, we converted National Marrow Donor Program (NMDP) high resolution allele frequencies to antigen equivalents and compared with the UNOS Calculated Panel Reactive Antibodies (CPRA) reference panel.

**Results:** Normalized frequency similarity scores between independent NMDP and UNOS panels for 4 US population categories (Caucasian, Hispanic, African American and Asian/Pacific Islander) ranged from 0.85 to 0.97, indicating correct antigen mapping. An open source web application (ALLele to ANtigen (“ALLAN”)) and web services were also developed to map unambiguous and ambiguous HLA typing data to UNOS antigen equivalents based on NMDP population-specific allele frequencies (http://www.transplanttoolbox.org).

**Conclusions:** This tool sets a foundation for using molecular HLA typing to compute the virtual crossmatch and may aid in reducing typing discrepancies in UNet.

## Introduction

Human leukocyte antigen (HLA) specificities were originally distinguished by allosera from multiparous women or individuals with multiple blood transfusions using cell-based testing strategies. HLA typing is currently performed by more accurate and specific molecular methods that assay nucleotide sequences. The IMGT/HLA database was developed to catalog the growing number of known HLA gene sequences^1^. While the naming of HLA sequences has its roots in the antigen nomenclature from serologic typing^2^, new HLA alleles are named based on sequence similarity to known alleles.

The United Network for Organ Sharing (UNOS) organ allocation system UNet utilizes donor and recipient HLA typing information and a list of unacceptable HLA antigens determined from antibody assays to perform virtual crossmatch as part of organ offers^3,4^. While the more specific IMGT/HLA allele nomenclature is used to interpret DNA-based typing data, only serologic antigen equivalents are accepted by the UNet^5^. Antigen groups are used to represent unacceptable HLA specificities because a single allo-HLA antibody may react similarly against several distinct HLA alleles within the same antigen group.

Histocompatibility labs face a data management challenge in mapping molecular-level HLA data to UNOS antigen equivalents for entry into UNet. While the Organ Procurement and Transplantation Network (OPTN) published a policy document in 2003 with general guidelines for mapping molecular data to antigen equivalents^6^, there is a need to automate the mapping process with a standardized mapping table that includes every IMGT/HLA allele. UNOS deemed it infeasible to maintain such a table at the time. Because of the difficulty in synthesizing antigen equivalency information from multiple reference data sources, there is likely to be some variability in reporting of HLA typing results into UNOS systems, even among commercial typing platforms.

A rapid turnaround for HLA typing is necessary for limiting cold ischemic time, therefore deceased donors are typed by sequence-specific primer (SSP), sequence-specific oligonucleotide probe (SSOP), or real-time polymerase chain reaction (PCR) based methods rather than next-generation sequencing (NGS). These methods rarely achieve allele-level specificity. Hence, there is also a need to interpret ambiguous HLA typings in the context of population HLA frequencies, as OPTN guidelines call for reporting UNOS antigens based on the most common HLA allele.

To meet these challenges in HLA data reporting for virtual crossmatch, we were inspired by matching systems for hematopoietic stem cell transplantation (HSCT) that have the capability to represent HLA typing data in IMGT/HLA allele nomenclature and match to donors whose HLA is represented in antigen nomenclature^7^. The World Health Organization (WHO) Nomenclature Committee for Factors of the HLA System maintains a list of official WHO antigen equivalencies for common HLA alleles. For HSCT registry matching, the IMGT/HLA database also maintains a World Marrow Donor Association (WMDA) antigen equivalency table^8^ that is updated quarterly with each database release^9^. No such mapping table exists for UNOS antigen equivalents. UNOS antigen equivalents differ from WHO and WMDA antigens for some alleles and we found these differences can be derived from interpreting UNOS/OPTN guidelines and using UNOS-provided tables that list IMGT/HLA alleles that are exceptions to these guidelines.

To improve fidelity in the communication of HLA data from histocompatibility labs to the UNOS allocation system, we describe a mapping table to convert IMGT/HLA alleles to UNOS antigen equivalents. We have also made available open source informatics tools to implement the aforementioned UNOS/OPTN guidelines, to keep the mapping table current with each IMGT/HLA database release, and to perform conversion of both ambiguous and unambiguous molecular HLA typing to UNOS antigens. We further describe how these tools will enable molecular HLA typing data to be used directly to compute the virtual crossmatch.

## Materials and Methods

### Data Sources

#### UNOS/OPTN Policy Documents for “Interpretation of HLA Typing Results for Entry into UNet”

We have summarized the rules for histocompatibility labs to assign UNOS antigens for IMGT/HLA alleles in Table 1. These rules were developed based on interpretation of guidelines published in 2003 in an OPTN policy document^6^. The OPTN documents^5,6^ contain tables of UNOS antigen equivalencies for some IMGT/HLA alleles that differ from WHO-assigned antigens. These tables include some 4-digit antigens representing alleles that may be readily identified by low resolution molecular testing as well as antigens that were not recognized by WHO serologic nomenclature at the time of policy implementation. The UNOS Histocompatibility Committee issued updates in 2017 that specified antigen assignments for some additional IMGT/HLA alleles^10^.

**Table 1:**
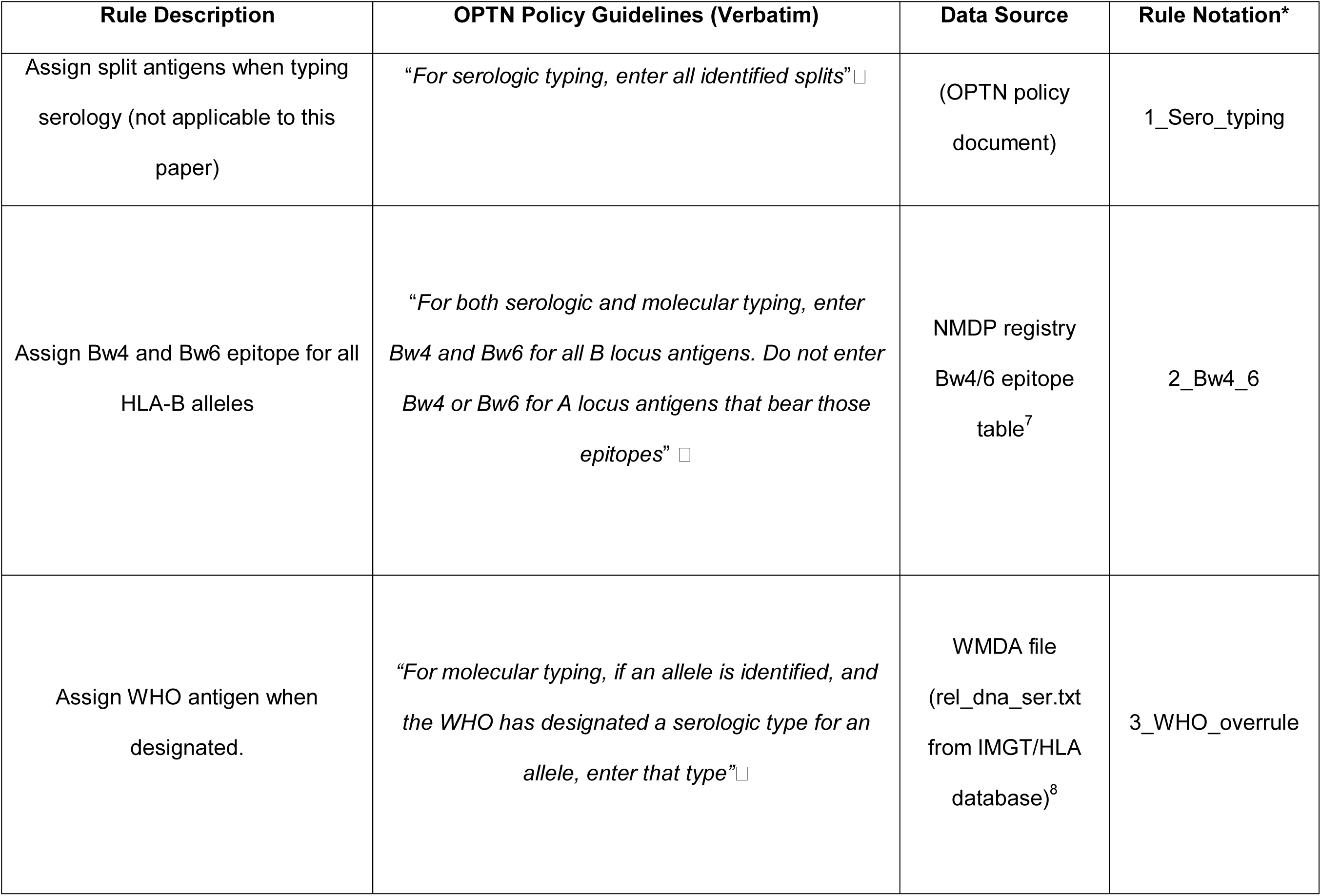
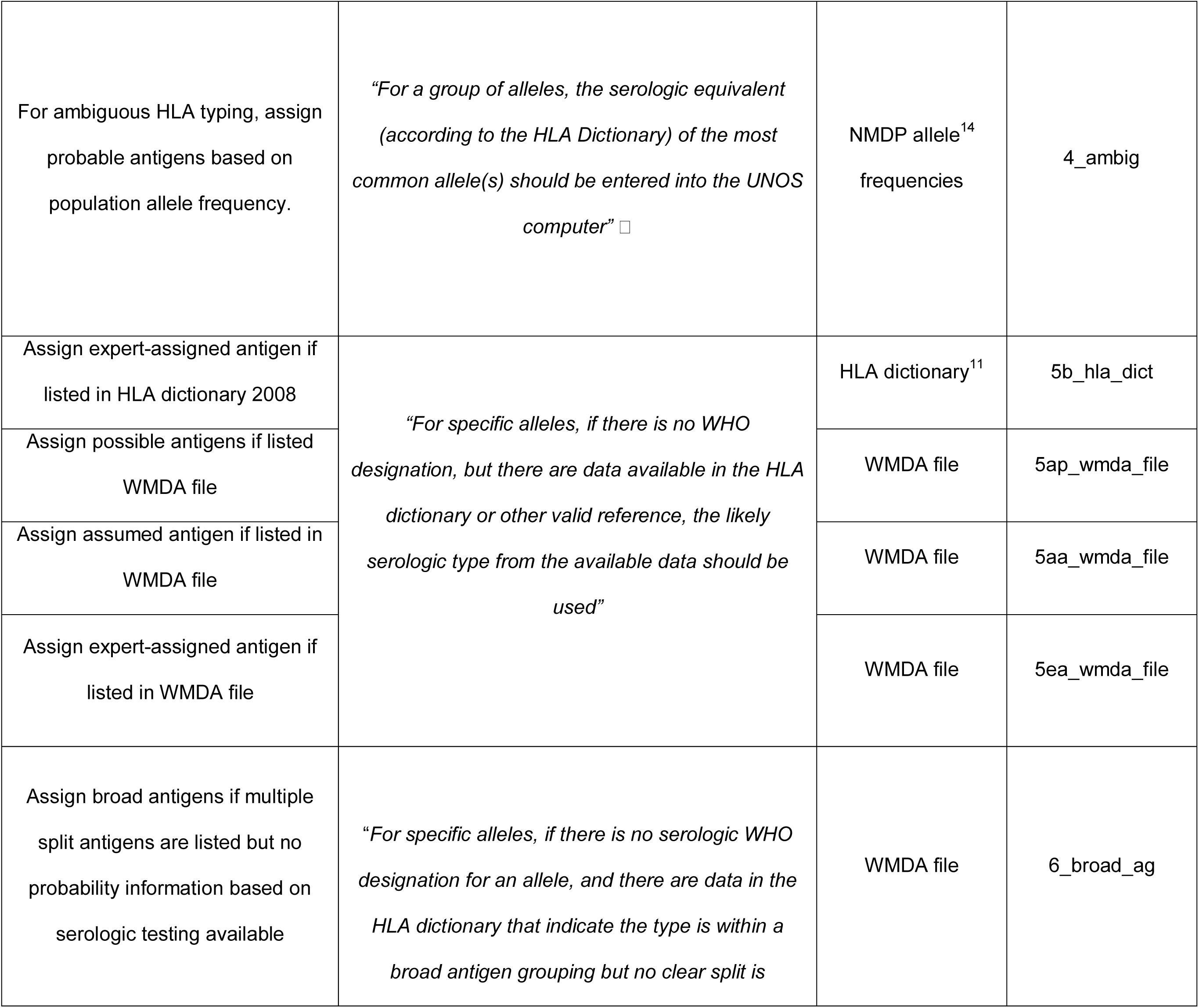
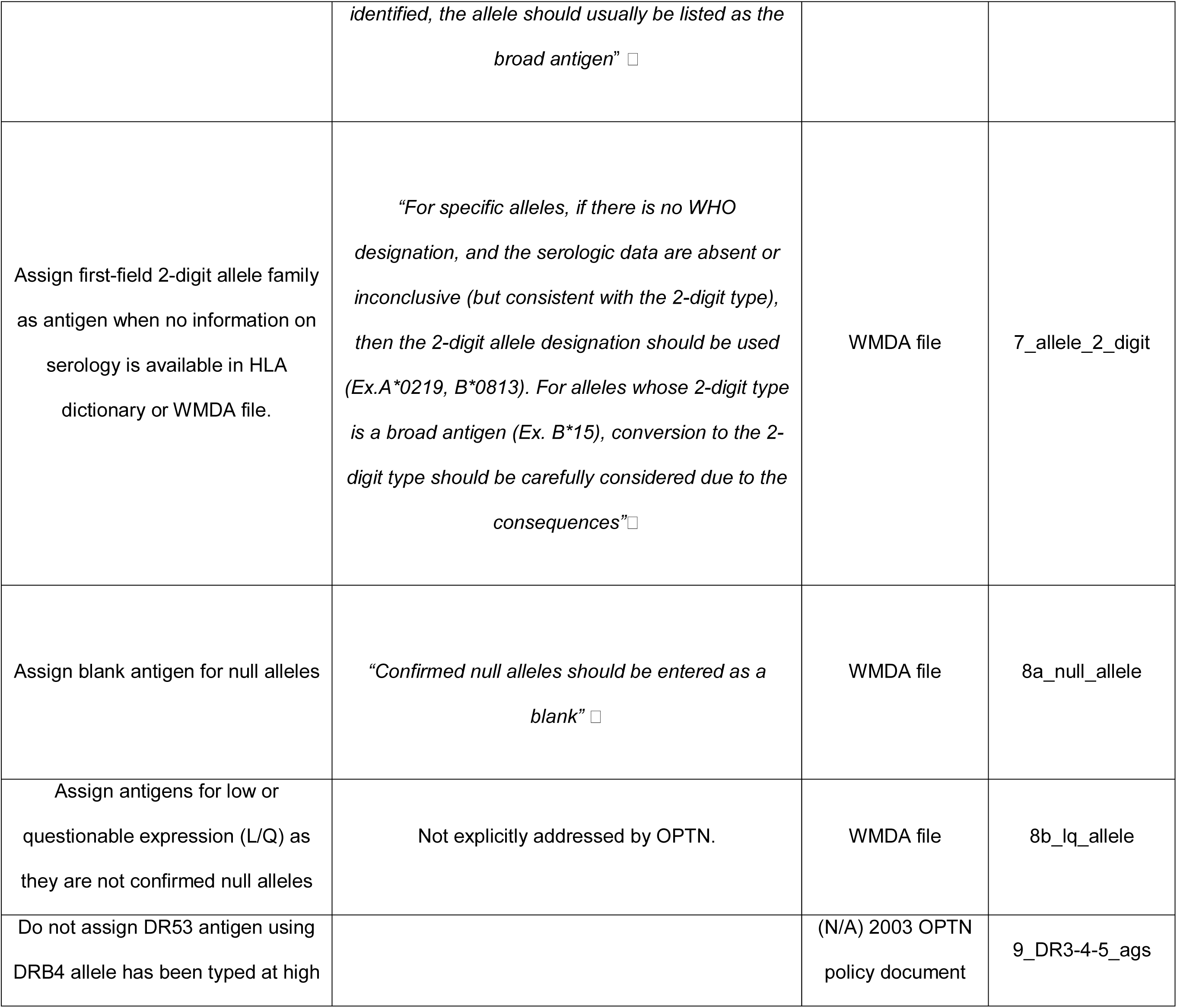
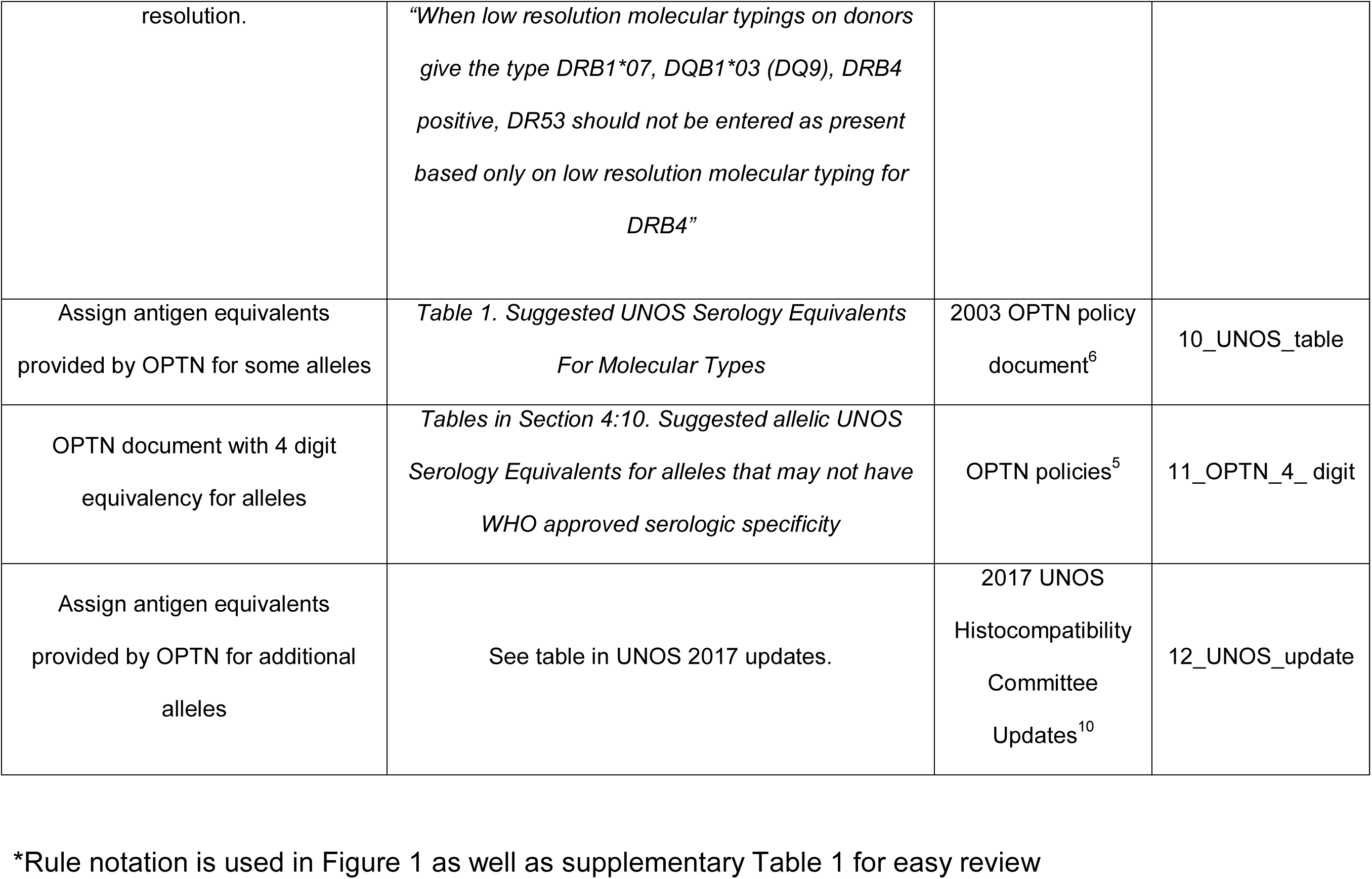
Rules for Mapping Molecular HLA typing to UNOS Antigen equivalents.

#### HLA Dictionary 2008

To standardize HLA antigen assignments for stem cell donor registries, the WMDA maintains an HLA Dictionary^11^. The 2008 version of the HLA Dictionary describes serologic equivalents of alleles for 6 loci (A, B, C, DRB1, DRB3/4/5, and DQB1). The data sources used to compile the dictionary include the following: WHO serological assignments as described in the Nomenclature report of 2004^12^, data submitted by individual laboratories to WHO committee for factors of HLA system, the international cell exchange program that has 200 laboratories participating in evaluating the standardized sera, NMDP-assigned serological equivalents, and data submitted for 13^th^ International HLA and Immunogenetics Workshop (IHIWS). Besides these sources that assigned serotypes based on the testing of cells with sera, the dictionary also lists antigens that were computationally assigned by a neural network^13^. Final consensus antigen assignments are listed as an expert-assigned type.

#### WHO and WMDA Antigen Equivalents Maintained by IMGT/HLA

The quarterly updated IMGT/HLA database curates the sequences of HLA alleles. The current release (v.3.32.0) includes 16,886 alleles for A, B, C, DRB1, DQA1, DQB1 and DRB3/4/5 loci. Each release includes a file with the WHO and WMDA antigen assignments for every IMGT/HLA allele^8^ (rel_ser_dna.txt, referred as the “WMDA File” in this paper). There are several categories of antigen equivalencies in this file based on the amount of information available from serologic testing. Many of the common IMGT/HLA alleles have WHO antigen assignments with unambiguous serology. WHO assignments were either made by the nomenclature committee when the allele sequence was submitted or were based on serologic testing data from the HLA Dictionary^11^. Possible serological antigens are given when there is no serologic testing data and are separated by a slash (“/”) when there is more than one possibility. Assumed antigens are typically based on first field of the IMGT/HLA allele name. WMDA expert-assigned antigens come from search determinants assigned by stem cell donor registries. UNOS antigen assignments for new IMGT/HLA alleles can thus rely upon updates to this WMDA file.

#### NMDP Allele Frequencies for Assigning Antigens for Ambiguous HLA typing

We used population-specific high resolution allele frequencies to calculate the frequencies of all possible UNOS antigens and provide the most probable antigen as the best assignment. High resolution allele frequencies for 4 broad US race/ethnic groups (Caucasian (CAU), African American (AFA), Hispanic (HIS), and Asian/Pacific Islander (API)) for the A, B, C, DRB3/4/5 DRB1, and DQB1 loci were obtained from NMDP^14^. These frequencies have been derived from 6.59 million volunteer donors from the Be The Match registry for HSCT. The NMDP US frequencies also include the Native American broad race and 21 detailed race/ethnic subcategories that could potentially be selected to make more precise antigen assignments.

## Implementation

### Conversion Table Program and Functions

The program to generate the IMGT/HLA allele to UNOS antigen conversion table was written in Python. The conversion table includes all current IMGT/HLA alleles at A, C, B, DRB1, DRB3/4/5, DQA1, DQB1 loci and their respective UNOS antigen equivalents according to UNOS guidelines shown in Table 1. A command line Python conversion script supports four different input forms of HLA typing results: unambiguous HLA alleles (either a single allele or a list of alleles) and ambiguous multi-locus HLA typing (represented as a genotype list string (GL string)^15^ or as NMDP multiple allele codes (MACs)^16^). For ambiguous HLA typings, a US population must be selected, as the most probable antigens are calculated given the population-specific allele frequencies and the antigen mappings for the alleles. An up-to-date list of MACs is accessed via the NMDP MAC web service (https://hml.nmdp.org/mac/). A function for reverse mapping of an unacceptable UNOS antigen to a list of IMGT/HLA alleles is also provided. DQA1 antigen mapping for ambiguous typing is not included in this release, as interpretation requires population-specific allele frequencies that are under development by NMDP.

### Command Line Tool, Web Application, and Web Services

We developed a web application named ALLele to ANtigen (“ALLAN”) for users to enter molecular HLA typings and perform UNOS antigen conversion. The tool is available athttp://www.transplanttoolbox.org. This web tool was implemented using the Django web framework^17^. As a programming interface, Representational State Transfer (RESTful)^18^ enabled web services available at http://www.transplanttoolbox.org/tool_services were developed using Django REST Framework^19^. An example Python script that illustrates how to access the web services is also provided in our GitHub source code repository (https://github.com/lgragert/hla-who-to-unos). For advanced users who wish to run the conversion tool locally, a Python command line tool and pip installable package “transplanttoolbox-allan” are also available (https://pypi.org/project/transplanttoolbox-allan/).

### Antigen Frequencies and Similarity Index Calculation

Reference antigen frequencies for 4 broad US race/ethnic categories (CAU, AFA, HIS, and API) were generated by mapping NMDP haplotype frequencies to UNOS antigen equivalencies. To assess accuracy in antigen assignments we used a normalized similarity index, “I_F_” ^20^, to compare the converted NMDP antigen frequencies to an independent antigen-level frequency dataset of previous UNOS donors^21^ that is currently used operationally as the reference panel for measuring CPRA values.

## Results

### Mapping Serological Specificities for IMGT/HLA Alleles Based on UNOS Guidelines

An algorithm was designed to map IMGT/HLA alleles to antigens following the UNOS guidelines. Our interpretation of the rule precedence for antigen mapping guidelines is shown as a schematic diagram in Figure 1. The antigen assignments found in tables from UNOS histocompatibility committee updates, UNOS antigen equivalency tables for specific alleles (Table 1: Suggested UNOS serology equivalents for molecular types (2003))^6^, and 4-digit antigens defined by UNOS were given the highest precedence^5^. For the next level of precedence, WHO-assigned antigens from the WMDA file were assigned (see Rule 3 in Table 1) followed by antigens from HLA Dictionary 2008^11^. No antigens are assigned for DQA1 locus alleles in the WMDA file, however, UNOS has added 2^nd^-field allele digits as DQA1 antigen equivalencies in their latest update (e.g. DQA01:01 for DQA1*01:01:01:01).

**Figure 1:**
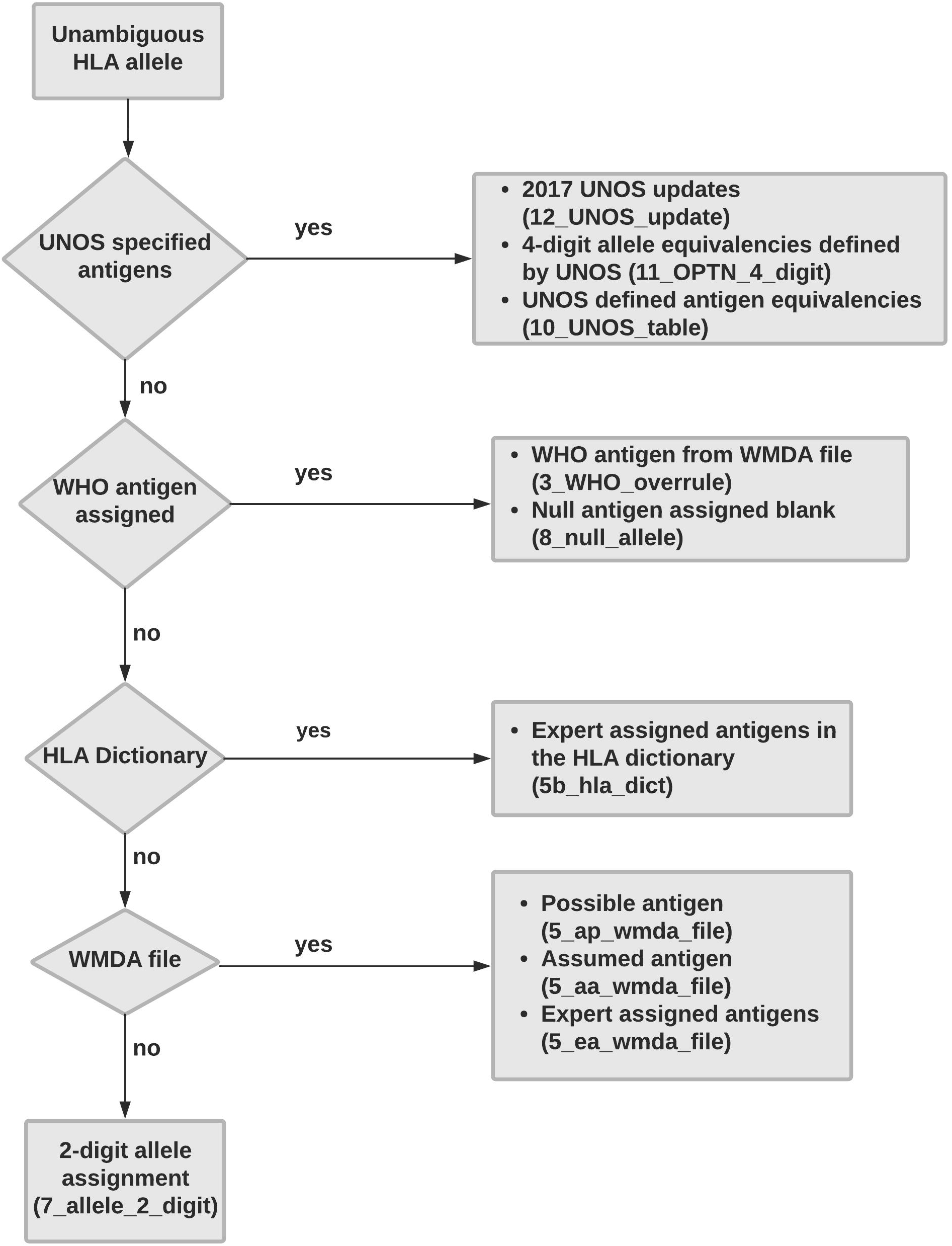
Flow diagram depicting defined rule precedence for UNOS histocompatibility committee guidelines. Highest precedence for antigen assignment to IMGT/HLA alleles was given to UNOS resources, followed by WHO antigens, the HLA dictionary, and the WMDA antigen file. If an antigen could not be assigned to an allele from these antigen equivalency reference sources, then the first-field of the IMGT/HLA allele name was assigned as the antigen.

If antigens could not be assigned from the data sources listed above, then possible and assumed antigens from the WMDA file were used. Cases where we fell back to assigning 2-digit allele equivalencies when information from all other data sources was lacking were mostly restricted to B and C locus alleles. When in doubt, we erred on the side of assigning a broader antigen category rather than assuming a split antigen.

A complete conversion table of UNOS antigen equivalencies for the IMGT/HLA alleles in release v.3.32.0 is available as Supplementary Table 1. We also provide a table that lists the IMGT/HLA alleles that correspond to an unacceptable antigen as Supplementary Table 2. Future conversion tables will be available on http://www.transplanttoolbox.org in tandem with quarterly updates to IMGT/HLA.

**Table 2:**
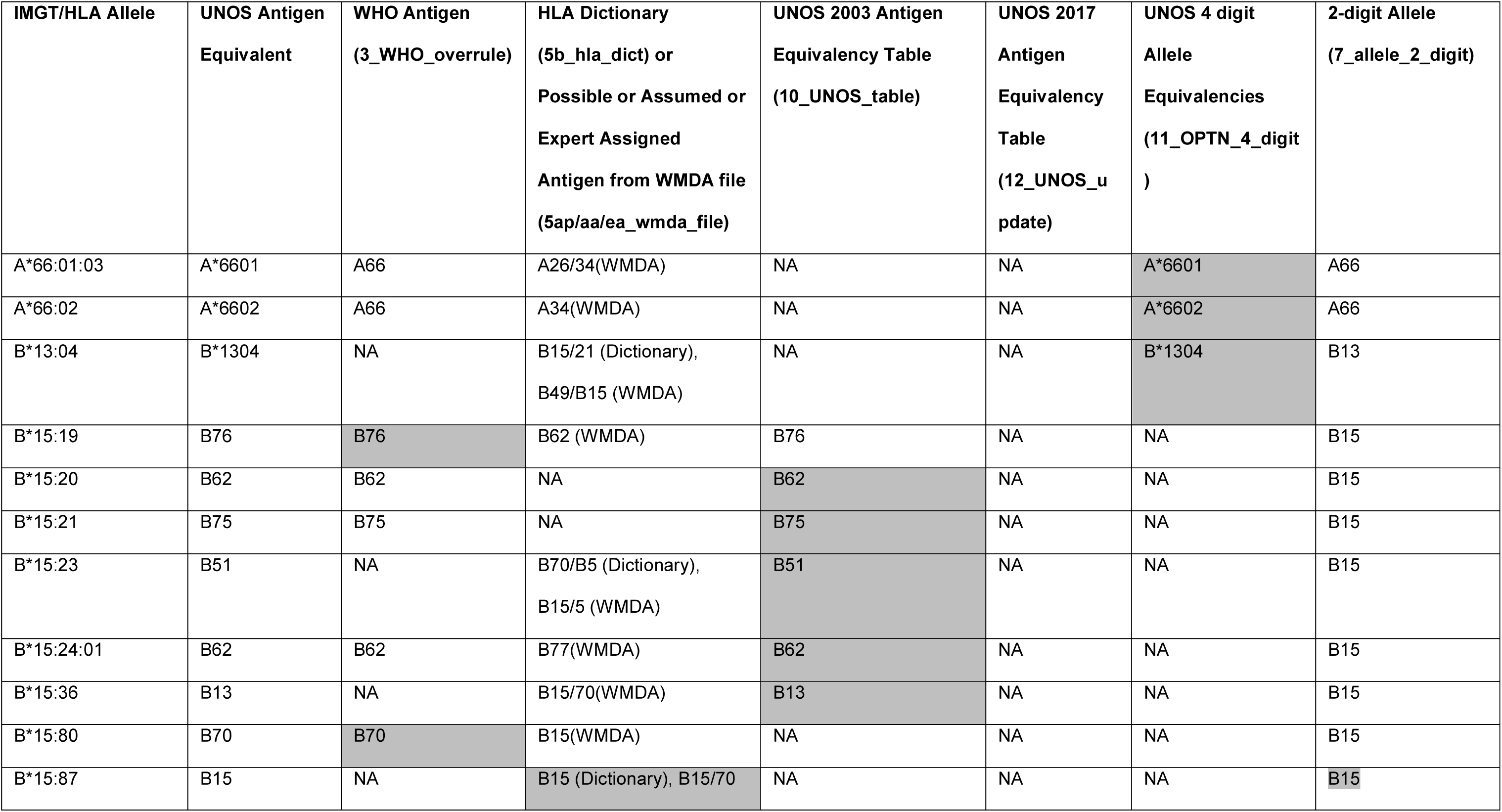
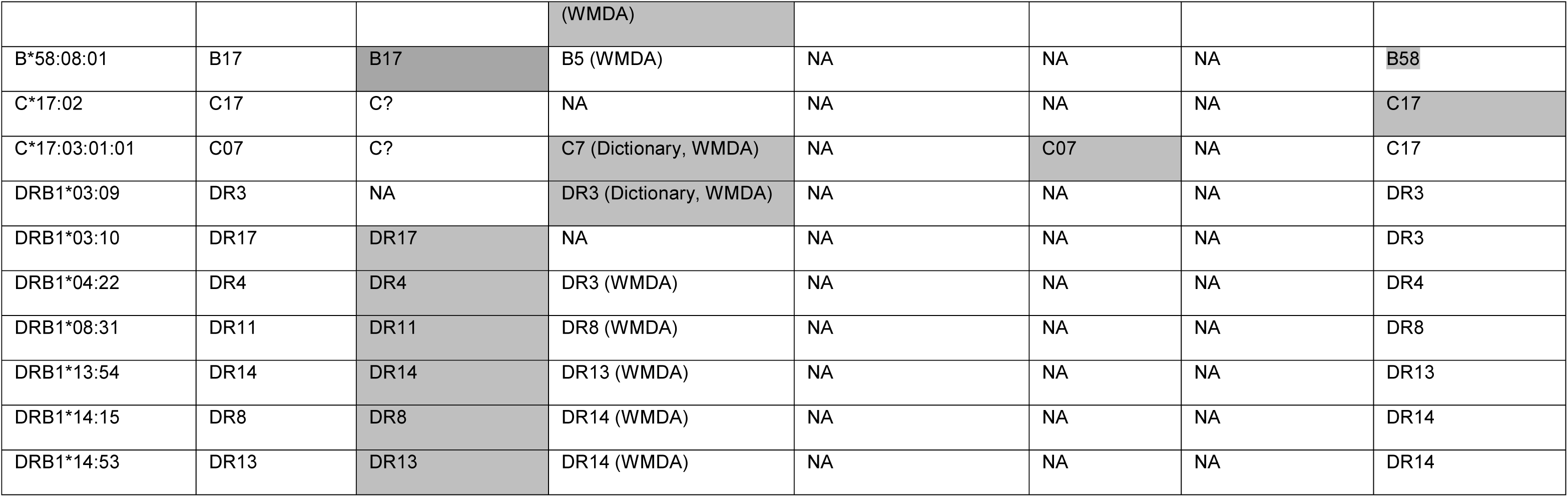
Illustration of Disparate Antigen Assignments Among Data Sources. The UNOS Antigen Equivalent column gives the final call given by the conversion tool. Disparities are resolved by the rule precedence in the flowchart in Figure 1. Shaded column indicates which data source was used to make the final call for few of the IMGT/HLA alleles.

### Precedence Order when Multiple Rules for Assigning Antigen Specificity may Apply

Multiple guidelines for antigen assignments may apply to a single IMGT/HLA allele, therefore an order of precedence for these guidelines must be set. When the UNOS antigen assignment listed in the UNOS-provided table conflicted with the WHO assignment, we chose the UNOS table. For example, for allele B*15:29, the WHO assignment is B15, whereas the UNOS assignment is B70.

It is not always correct to simply reduce allele nomenclature to 2-digits to get antigen specificity because different alleles from the same allele group can have different antigens. This is most evident in the B*15 group, where alleles may have antigen equivalents of B15, B62, B63, B70, B71, B72, B75, B76, or B77. Rule precedence resolves cases where different data sources suggest differing antigen assignments, as we illustrate with several examples in Table 2.

### Most Probable Antigen for Ambiguous HLA Typing

As per OPTN guidelines, for a group of possible alleles in an ambiguous HLA typing, the serologic equivalent of the most common antigen should be entered. We apply reference population-specific allele frequencies from the NMDP population categories to make the assignment. We show that for a particular ambiguous HLA typing, the most probable antigen for the B locus may vary depending on the broad race/ethnic category selected (Table 3).

**Table 3:** Antigen Mapping for Unambiguous and Ambiguous HLA Typing by The Conversion Tool “ALLAN”.

| HLA typing data input |  | Input example | Population selection | Antigen assignment by tool (antigen probabilities mentioned in parenthesis) |
| --- | --- | --- | --- | --- |
| Unambiguous typing | Allele | B*13:01:01:01 | Not required | B13, Bw4 |
|  | Allele Lists | A*02:301N B*42:08 C*03:04:01:08,<br>DQB1*03:241<br>DRB5*01:07 | Not required | A0, B42, C10, DQ3,<br>DR51, Bw6 |
| Ambiguous typing | Genotype List String | A*02:01+A*26:01g^C*05:01g+C*01:02g<br>^B*15:01/B*15:02/B*15:03/B*15:04+B*39:01/B*39:02/B*39:04/B*39:05 | African American | A2 + A26 (1.0),<br>C05 + C01 (1.0),<br>B72 + B3901 (0.77),<br>Bw6 |
|  |  |  | Asian/Pacific Islander | A2 + A26 (1.0),<br>C05 + C01 (1.0),<br>B75 + B3901 (0.52),<br>Bw6 |
|  |  |  | Caucasian | A2 + A26 (1.0),<br>C05 + C01 (1.0),<br>B62 + B3901 (0.96),<br>Bw6 |
|  |  |  | Hispanic | A2 + A26 (1.0),<br>C05 + C01 (1.0),<br>B62 + B39 (0.47),<br>Bw6 |
|  | NMDP<br>Multiple<br>Allele Codes | A*01:AABJE A*02:HBMC B*08:NMTJ<br>B*13:GR C*04:CYMD C*05:YDYE<br>DRB1*04:AMR DRB1*07:GC<br>DQB1*03:AG DQB1*03:AFYYJ | African<br>American | A1 + A2 (0.50),<br>C04 + C05 (0.99),<br>B8 + B13 (1.0),<br>DR4 + DR7 (1.0),<br>DQ7 + DQ3 (0.82),<br>Bw4, Bw6 |
|  |  |  | Asian/Pacific<br>Islander | A1 + A2 (0.50),<br>C04 + C05 (0.90),<br>B8 + B13 (1.0),<br>DR4 + DR7 (1.0),<br>DQ7 + DQ3 (0.82),<br>Bw4, Bw6 |
|  |  |  | Caucasian | A1 + A2 (0.49),<br>C04 + C05 (0.99),<br>B8 + B13 (1.0),<br>DR4 + DR7 (1.0),<br>DQ7 + DQ3 (0.85),<br>Bw4, Bw6 |
|  |  |  | Hispanic | A1 + A2 (0.80),<br>C04 + C05 (0.99)<br>B8 + B13 (1.0),<br>DR4 + DR7 (1.0),<br>DQ7 + DQ3 (0.82),<br>Bw4, Bw6 |

### “ALLAN”: IMGT/HLA ALLele to UNOS ANtigen conversion tool

The conversion functions described above are available via the ALLAN web tool at http://www.transplanttoolbox.org. When the HLA typing is unambiguous, users may select to enter either a single allele or a list of alleles. When the HLA typing contains typing ambiguity, users may enter the typing either in GL string or MAC format (Table 3). The output shows the UNOS antigen assignments and Bw4/6 epitopes. Entry of ambiguous HLA typings requires the selection of a US race/ethnic group and returns the probability distribution of all the possible antigens considering the allele frequencies in the selected race/ethnic group. A complete user guide for the tool is available on the website. For advanced users, we also developed a Python script as a command line tool with similar functionality, which can be accessed via our GitHub repository (https://github.com/lgragert//hla-who-to-unos). We have also developed web services as a programming interface for researchers and laboratory information management systems. Client applications such as HLA typing software platforms or other lab information systems can send HLA typing and race/ethnicity information to the service, and UNOS antigens (including probabilities) are returned. The endpoint URLs and commands for these services are shown in Table 4.

**Table 4:**
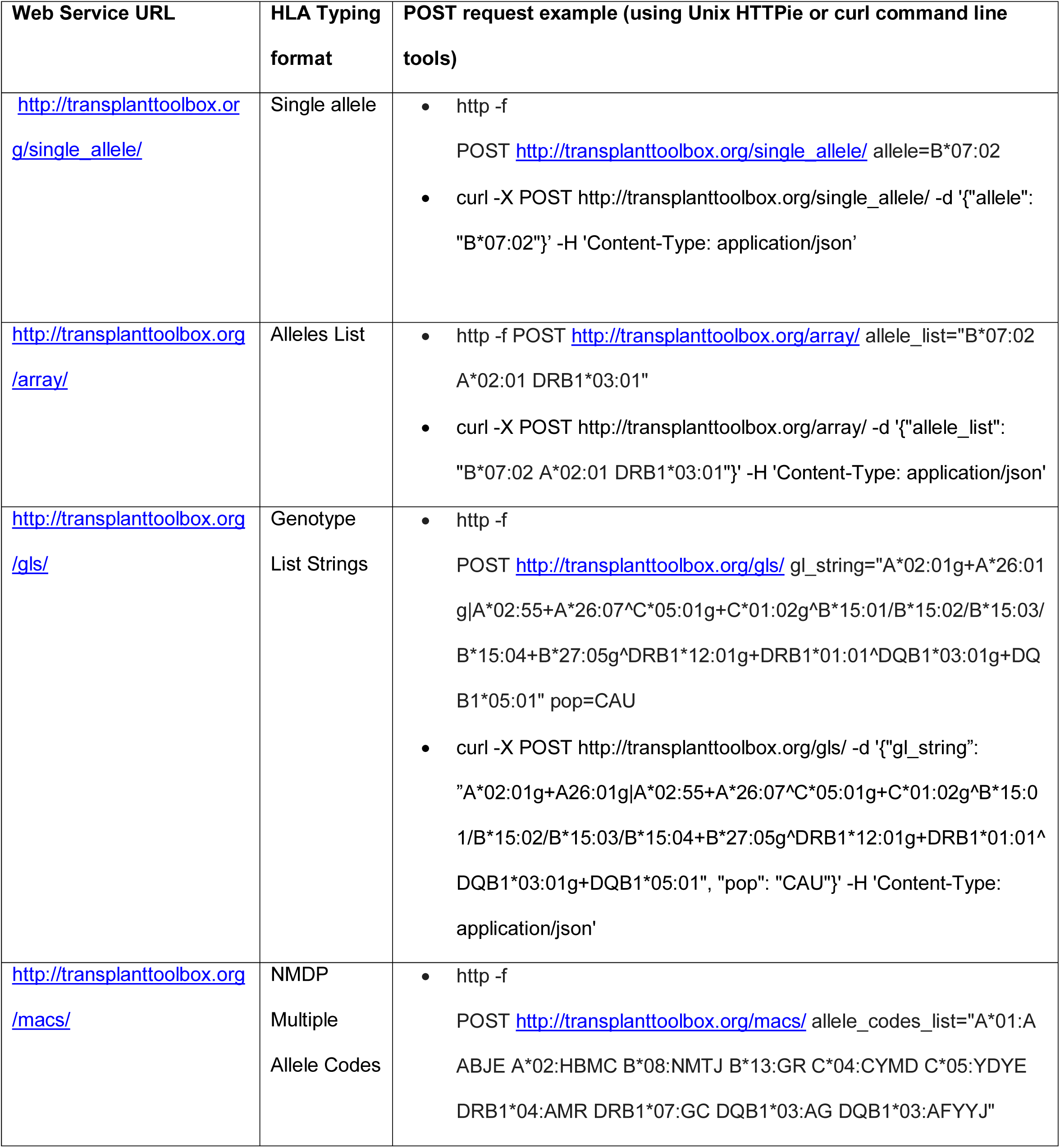
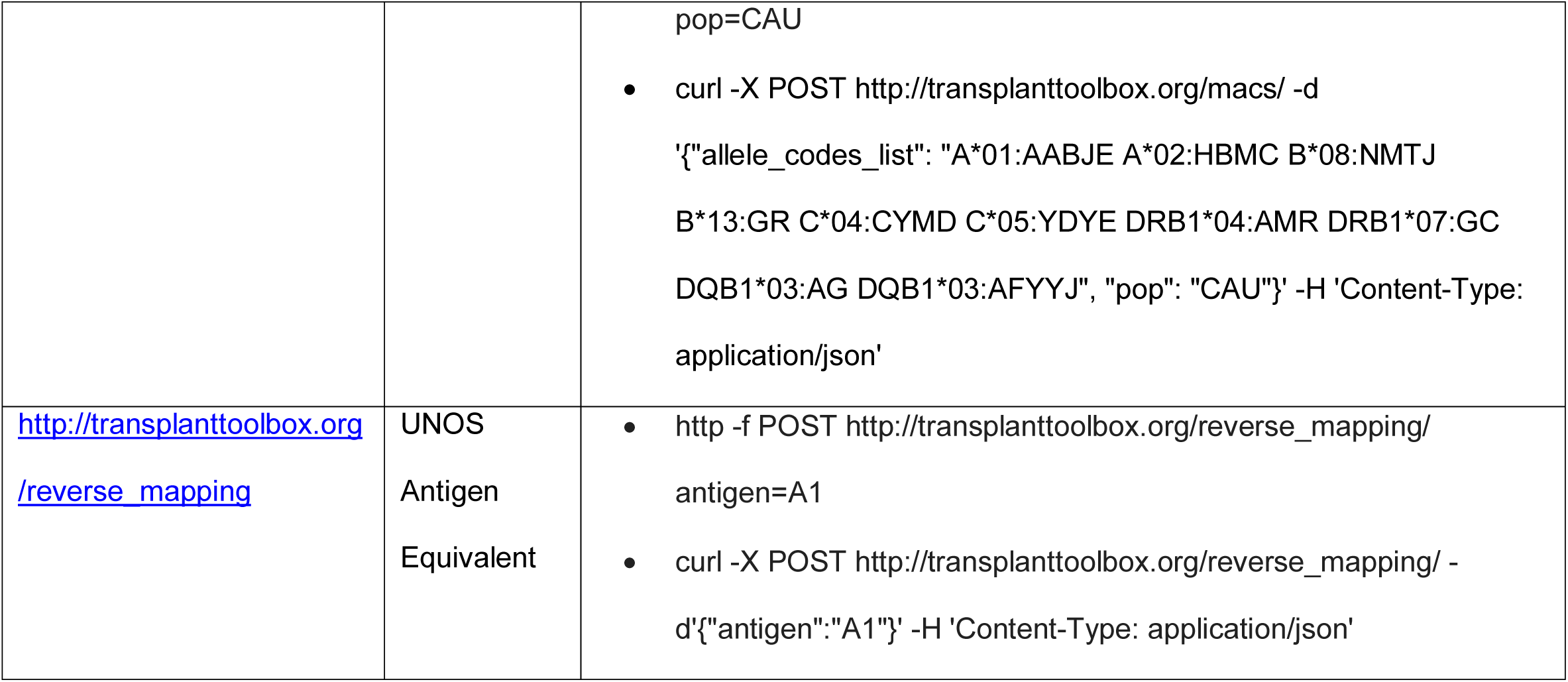
“ALLAN” Web Services for UNOS Antigen Assignments and Reverse Mapping.

### Antigen Mappings in Commercially Available Software Programs

Most histocompatibility labs rely on commercial software to analyze HLA typing results and perform antigen conversion. Currently, the software platforms from leading vendors in this domain, i.e. One Lambda (HLA Fusion) and Immucor (LIFECODES^®^ MATCH IT!^®^), perform antigen mapping for a limited number of IMGT/HLA alleles but would benefit from a complete standardized table. “HLA Fusion” does mapping for 4,927 alleles and “MATCH IT!” does mapping for 4,439 of the 16,886 alleles. Compliance with IMGT database updates for new releases of HLA alleles and UNOS updates represents a major maintenance burden for these software vendors. Vendors must rely on available sources such as the HLA Dictionary, which has not been updated since 2008. The WMDA does provide quarterly updates, but again vendors are left with choosing between “Unambiguous associations,” “Possible associations,” “Assumed associations,” and “Expert assigned exceptions,” and have to reconcile these options with customer specific requests. As a result, some antigen mappings made by these software packages differ from those recommended by UNOS. A list of discrepancies from apparent UNOS guidelines, apart from the missing mappings, is shown in Table 5. The commercial software vendors are in agreement that the issues arising from independent maintenance and interpretation of antigen conversion rules will be alleviated by having a consensus set of mappings.

**Table 5:** Discrepancies in UNOS Antigen Assignments between the Conversion Tool and Commercial HLA Typing Software Packages.

| **Allele** | **UNOS Antigen**<br>**Equivalency**<br>**From ALLAN** | **One**<br>**Lambda**<br>**Fusion** | **Immucor**<br>**Lifecodes** |
| --- | --- | --- | --- |
| A*02:10<sup>a</sup> | A2 | A210 | A210 |
| A*66:01:01:01 | A*6601 | A66 | A66 |
| A*66:01:01:02 | A*6601 | A66 | A66 |
| A*66:01:02 | A*6601 | A66 | A66 |
| A*66:01:03 | A*6601 | A66 | A66 |
| A*66:02 | A*6602 | A66 | A66 |
| B*07:03<sup>a</sup> | B7 | B703 | B703 |
| B*15:29 | B70 | B15 | B15 |
| B*15:46 | B50 | B72 | B72 |
| B*15:58 | B62 | B62 | B15 |
| B*27:08 | B2708 | B2708 | B27 |
| B*35:76 | B35 | B35/22 | B35/22 |
| B*39:01:01:02L | B3901 | B39"Low" | Low B3901 |
| B*47:02 | B60 | B47 | B47 |
| B*51:03 | B51 | B5103 | B5103 |
| B*82:01 | B*8201 | B82 | B82 |

<sup>a</sup>New antigen equivalency provided by 2017 UNOS updates

### Reference Antigen Frequencies for Six HLA Loci in 4 Broad US Races

To validate the antigen assignments, we converted published NMDP high resolution haplotype frequencies^14^ into antigen equivalents and compared antigen frequencies at each HLA locus to the UNOS CPRA HLA reference panel. Antigen-level NMDP frequency tables for 4 US broad race/ethnic categories are available as Supplementary Table 3. The normalized “I_F_” similarity index for these races ranged from 0.85 to 0.97, indicating accuracy in antigen assignments (Table 6). One cause of disparity between panels was that the UNOS DQ typing data was more often reported as broad DQ1 antigens rather than split DQ5 and DQ6 antigens. Meanwhile, the only high resolution IMGT/HLA alleles in the NMDP panel that mapped to broad DQ1 were the relatively uncommon alleles DQB1*06:11 and DQB1*06:12.

**Table 6:** Similarity between UNOS Antigen Frequencies and NMDP High Resolution Allele Frequencies after Conversion to Antigen Equivalents. Normalized similarity index “I_F_” values compared NMDP-derived antigen frequencies to the UNOS CPRA reference panel for A, B, C, DR, DQ loci for four broad US race/ethnic categories: Caucasian, African American, Hispanic and Asian/Pacific Islander.

| **Locus** | **Caucasian** | **African**<br>**American** | **Hispanic** | **Asian/Pacific**<br>**Islander** |
| --- | --- | --- | --- | --- |
| A | 0.97 | 0.95 | 0.94 | 0.87 |
| B | 0.93 | 0.91 | 0.88 | 0.85 |
| C | 0.90 | 0.86 | 0.87 | 0.86 |
| DR | 0.96 | 0.96 | 0.95 | 0.88 |
| DQ | 0.89 | 0.90 | 0.90 | 0.89 |

## Discussion

The OPTN guidelines for HLA allele to antigen mapping require that histocompatibility labs refer to several different data sources which as we have shown here can lead to differences in antigen assignment. To streamline this process, we present a complete mapping table between all current IMGT/HLA alleles and UNOS antigens that was developed based on published OPTN guidelines. This conversion table report gives a standardized source for antigen mapping that can be updated in tandem with every new IMGT/HLA database release.

Our web tool, ALLAN, offers a convenient mapping of HLA typing data to UNOS antigens for entry into UNet, including resolving HLA typing ambiguities in antigen assignments as shown in Table 3. This will aid in streamlining HLA typing that is performed for deceased donors, which has limited time for experimentally resolving ambiguities. Molecular methods such as SSP may not always resolve null expression variants in time, however, our tool helps in predicting the probability of null alleles and therefore a blank antigen for a specified population.

One potential cause of unexpected positive tissue crossmatch^22^ post organ shipment is inaccuracy in manual HLA data entry, as the accuracy of prediction of negative virtual crossmatch relies solely on HLA data mapped and entered. A report from 22 organ procurement organizations (OPOs) describing the allocation practices of kidneys in high PRA patients indicates that a substantial number of kidneys shipped were transplanted in unintended recipients (19.2%) or discarded (3.5%) because of a positive crossmatch^23^. An analysis by the Discrepant HLA Typing Subcommittee of the UNOS Histocompatibility Committee showed that among 18,719 deceased donors whose HLA data were entered into UNet from 2015-2016, 6,670 (35.6%) had some discrepancy in their typing including allele vs antigen resolution, broad/split antigen nomenclature differences, or incorrect Bw4/6 or DRB3/4/5 assignments. 2% had a critical antigen-level discrepancy that would have affected the match run or allocation (Personal communication – Peter Lalli ^24^). Automated management of HLA typing data, including the use of a standardized antigen mapping table or direct use of molecular typing data to compute the virtual crossmatch, has the potential of reducing such HLA discrepancies and the need for repeat match runs, thus reducing the cold ischemia time. Further improvement in accuracy of typing data in UNet could be achieved via adoption of the Histoimmunogenetics Markup Language (HML) data standard that is used today for automated transfer of HLA typing data from laboratories into the NMDP stem cell donor registry^25^.

Out of 16,886 current IMGT/HLA alleles mapped to UNOS antigen equivalents, 12,489 had clear assignments where only one of the OPTN mapping guidelines would apply, while the remaining required us to interpret rule precedence. The high similarity between independent UNOS and NMDP derived antigen frequencies increases confidence in the accuracy of the mapping table. A lower similarity between API samples compared to other populations is likely due to a shallow sampling of only 333 Asian individuals in the UNOS data.

In recent updates, UNOS increased the number of 4-digit unacceptable antigens that can be reported. Because many of these 4-digit antigens are not included in the UNOS antigen frequencies for CPRA^26^ and do not increase CPRA values when selected as unacceptable antigens, we chose to map to broader antigens to ensure compatibility with the current UNOS system.

Bw4/Bw6 epitope categories are determined based on amino acid sequence for positions 77 through 83 of HLA-B, and there has not been serologic testing of some of the rare sequence motifs. The OPTN policy document^6^ does not include assignments for all Bw4/Bw6-defining motifs among IMGT/HLA alleles, therefore we used NMDP assignments, some of which were manually curated^7^. For the rare Bw4/Bw6 motifs, there were some discrepancies with the commercial HLA typing software programs. This highlights the need for an improved curation process that would lead to the availability of public Bw4 and Bw6 epitope assignment resources.

Few low (L) or questionable (Q) expression alleles were assigned antigens in the HLA dictionary and we have assigned antigens to all these alleles as if they were expressed. While L alleles have been confirmed to have weak surface expression that can elicit alloimmunity, little information is available for Q alleles^27^. The current version of IMGT/HLA database includes 75 Q alleles, yet only 8 of these have been observed unambiguously in the NMDP registry donor file. Further examination of mRNA transcripts, cell surface expression, and tests of alloimmunity are needed to make definitive assignments for these alleles, but we prefer to err on the side of caution. The assigning of antigens to L/Q alleles intends to avoid cases of false negative virtual crossmatch at a potential cost of increasing positive virtual crossmatch predictions.

OPTN guidelines do not call for histocompatibility laboratories to analyze HLA sequences. Therefore, decisions on antigen assignments are best maintained in the reference data sources such as the IMGT/HLA database, and the HLA dictionary. These resources synthesize information from individual reports, such as describing B64, B65 antigen splits of the broad B14 antigen defined by Street and Darke^28,29^ for B*14:01 and B*14:02. While B*14:03 had a shared amino acid motif with B*14:02 that could define the epitope, B*14:03 could be deemed a B65 based on sequence analysis. However, both WHO and the HLA dictionary call B*14:03 a B14. As another example, in a recent report^30^ these authors describe the serological specificity of B*15:33 as a B62. OPTN guidelines call for the WHO assignment, which is B15^31^. If the allele lacked a WHO assignment, the 2008 HLA dictionary would be used to assign a B62. For the UNOS assignment to become B62, the WHO antigen would need to change or UNOS would need to specify an exception to the rule. In summary, these complex cases cannot all be sorted out here. We believe such decisions are best left to UNOS, WHO, or the next HLA Dictionary.

Standardized antigen assignments for molecular typing are necessary for the UNOS organ allocation system to evolve to incorporate DNA-based HLA assignments and epitope-based matching strategies^32^. We anticipate future updates to our tool to support changes in the UNOS CPRA to include antigens for the DQA1, DPA1, and DPB1 loci. These loci together account for >60% of the unacceptable antigens in candidates with CPRA > 99%^33,34^. We intend to develop additional tools that would allow the NMDP high resolution frequency panel to be used for allele-level CPRA calculations incorporating these additional loci. Imputation tools based on NMDP frequency data are also useful for estimating amino acid assignments from antigen-level data to perform epitope analysis with tools such as PIRCHE: predicted indirectly recognizable HLA epitopes^35^. The antigen mapping tools we developed here will take UNOS one step closer to their long-term goal of maximally utilizing HLA molecular typing data in solid organ allocation systems.

## Acknowledgments

We thank Anat Tambur and Malek Kamoun for their advice and review of early drafts of this manuscript. We also thank the UNOS Histocompatibility Committee for their support of this project.

## Supplemental digital content captions

**Supplementary Table 1:** Reference table for HLA antigen mapping as per UNOS guidelines for 16,886 IMGT/HLA alleles encompassing A, C, B, DRB1, DRB3/4/5, DQA1 and DQB1 loci.

**Supplementary Table 2:** Reference table for the list of IMGT/HLA alleles that correspond to each unacceptable UNOS antigen. The assignments utilize the unacceptable antigen equivalency tables specified in the OPTN Policies Section 4.10 - Reference Tables of HLA Antigen Values and Split Equivalences, Tables 4-5 through 4-11.

**Supplementary Table 3:** NMDP antigen-level HLA frequencies for the A, B, C, DR, and DQ loci for 4 US broad ethnic groups: Caucasian, African American, Hispanic and Asian/Pacific Islander. Frequency data was converted from previously published high resolution allele frequencies.

Authorship
Navchetan Kaur: Participated in research, analysis and writing of the paper
Evan P Kransdorf: Participated in research design, analysis and writing of the paper
Marcelo J Pando: Participated in research design, analysis and writing of the paper
Martin Maiers: Contributed the data sources used in the research
Bryan Ray: Contributed the data sources used in the research
Jar-How Lee: Contributed the data sources used in the research
Peter Lalli: Contributed the data sources used in the research
Robert A Bray: Contributed the data sources used in the research
Cathi L Murphey: Contributed the data sources used in the research
Loren Gragert: Participated in research design, performance of research, analysis and writing of the paper.

## Abbreviations

AFA: African American
ALLAN: ALLele to ANtigen
API: Asia / Pacific Islander
CAU: Caucasians
CPRA: calculated panel reactive antibodies
GL: genotype list
HIS: Hispanics
HLA: human leukocyte antigen
HML: Histoimmunogenetics Markup Language
HSCT: hematopoietic stem cell transplantation
IHIWS: International HLA and Immunogenetics Workshop
L/Q: Low expression or Questionable expression alleles
MAC: multiple allele code
NGS: next-generation sequencing
NMDP: National Marrow Donor Program
OPO: organ procurement organization
OPTN: Organ Procurement and Transplantation Network
PCR: polymerase chain reaction
REST: representational state transfer
SSOP: sequence-specific oligonucleotide probes
SSP: sequence-specific primers
UNOS: United Network for Organ Sharing
WHO: World Health Organization
WMDA: World Marrow Donor Association

